# Mechanism of µ-Opioid Receptor-Magnesium Interaction and Positive Allosteric Modulation

**DOI:** 10.1101/689612

**Authors:** X. Hu, D. Provasi, M. Filizola

**Affiliations:** Icahn School of Medicine at Mount Sinai

**Author notes:** Equal contributors (X. Hu, D. Provasi).

## Abstract

In the era of opioid abuse epidemics, there is an increased demand for understanding how opioid receptors can be allosterically modulated to guide the development of more effective and safer opioid therapies. Among the modulators of the µ-opioid (MOP) receptor, which is the pharmacological target for the majority of clinically used opioid drugs, are monovalent and divalent cations. Specifically, the monovalent sodium cation (Na^+^) has been known for decades to affect MOP receptor signaling by reducing agonist binding, whereas the divalent magnesium cation (Mg^2+^) has been shown to have the opposite effect, notwithstanding the presence of sodium chloride. While ultra-high resolution opioid receptor crystal structures have revealed a specific Na^+^ binding site and molecular dynamics (MD) simulation studies have supported the idea that this monovalent ion reduces agonist binding by stabilizing the receptor inactive state, the putative binding site of Mg^2+^ on the MOP receptor, as well as the molecular determinants responsible for its positive allosteric modulation of the receptor, are unknown. In this work, we carried out tens of microseconds of all-atom MD simulations to investigate the simultaneous binding of Mg^2+^ and Na^+^ cations to inactive and active crystal structures of the MOP receptor embedded in an explicit lipid/water environment. Analyses of these simulations shed light on (a) the preferred binding sites of Mg^2+^ on the MOP receptor, (b) details of the competition between Mg^2+^ and Na^+^ cations for specific sites, (c) estimates of binding affinities, and (d) testable hypotheses of the molecular mechanism underlying the positive allosteric modulation of the MOP receptor by the Mg^2+^ cation.

**Statement of Significance:** Overprescription of opioid drugs in the late 1990s, followed by abuse of both prescription and illicit opioid drugs, has led to what is nowadays called “opioid overdose crisis” or “opioid epidemic”, with more than 130 Americans dying daily from opioid overdose at the time of writing. To reduce opioid doses, and thereby prevent or treat overdose and opioid use disorders, attention has recently shifted to the use of co-analgesics. Understanding how opioid receptor targets can be allosterically modulated by these elements, including cations, is key to the development of improved therapeutics. Here, we provide an atomic-level understanding of the mechanism by which magnesium binds to the µ-opioid receptor and enhances opioid drug efficacy by stabilizing the receptor activated state.

## Introduction

Opioid analgesics such as morphine remain the “gold standard” for the treatment of acute (e.g., postoperative) or chronic (e.g., cancer) pain notwithstanding the multitude of side effects they cause, including respiratory depression, tolerance, addiction, dependence, constipation, nausea, vomiting, dizziness, fatigue, and itching (1). Among all side effects, respiratory depression is certainly the most feared by physicians given the almost 400,000 casualties from overdosing on prescription or illicit opioids during 1999-2017 in the United States alone (2).

Almost all clinically used opioid drugs act through the μ-opioid (MOP) receptor (3), a member of the G protein-coupled receptor (GPCR) superfamily. Like other GPCRs, this receptor exists as an ensemble of multiple inactive, active, and intermediate conformations, and this conformational plasticity forms the basis for the receptor signaling diversity achieved through receptor interaction with different intracellular proteins, such as G proteins (typically Gα_i/o_ subtypes) or arrestins (typically β-arrestin2) (4). Depending on the MOP receptor conformation stabilized by the drug and the intracellular protein recruited by the receptor, biological functions that are either clinically desirable or detrimental can ensue. Indeed, while studies using MOP receptor knockout mice confirmed that the MOP receptor is responsible for the anti-nociceptive action of morphine (3), β-arrestin recruitment by the MOP receptor was shown to contribute to some of the side effects of this classical opioid drug (5–7), supporting the hypothesis that biasing the conformational equilibrium of MOP receptor towards the G protein pathway may lead the way to develop safer opioid therapies.

In principle, there are several ways one could possibly bias the conformational equilibrium of the MOP receptor towards a specific conformational state, whether using a small molecule, a peptide, a protein, or other allosteric modulators, including cations. Indeed, it has been known for more than forty years that the MOP receptor can be differentially modulated by cations (8). While the monovalent Na^+^ cation can decrease agonist affinity at the MOP receptor (8), most likely though stabilization of the inactive conformational state of the receptor (e.g., see (9, 10)), the divalent Mg^2+^ cation has the opposite effect (e.g., see (11)), suggesting it stabilizes an active-like conformation of the receptor. Notably, similar conclusions were drawn for other GPCRs based on inferences from biochemical and pharmacological studies (12–18).

While various high-resolution X-ray crystal structures of inactive forms of GPCRs have revealed the atomic details of Na^+^ binding (19–22), and several MD simulation studies have supported the stabilization of inactive conformations of the receptor by this monovalent cation (e.g., see (9, 10, 23, 24)), limited information exists to date about the preferred binding site(s) of Mg^2+^ and the molecular mechanism underlying the potential positive allosteric effect of this divalent cation. A recent interdisciplinary study combining the results of ^19^F-NMR and MD simulations on another prototypic GPCR, the adenosine A_2A_ receptor (18), provided further support to the negative allosteric modulation of receptors by Na^+^ and their positive allosteric modulation by Ca^2+^ and Mg^2+^. Not only did this work allow to quantify the effects of these cations on the conformational ensemble of the adenosine A_2A_ receptor, but it also suggested important molecular determinants involved in the allosteric activation of this receptor in the presence of physiological cations. Notable results of these studies were the unexpected finding that Na^+^ also stabilized an intermediate state that had previously been associated to partial agonism, while Mg^2+^ cations drove G-protein-binding cleft opening, and consequent receptor activation, upon bridging specific acidic residues on the extracellular region of transmembrane helices 5 and 6 (TM5 and TM6) of the receptor (18).

In the present work, we studied the free binding of Mg^2+^ to the MOP receptor in the presence of physiological concentrations of Na^+^ by all-atom standard MD simulations of the receptor embedded in an explicit 1-palmitoyl-2-oleoyl-sn-glycero-3-phosphocholine (POPC)/cholesterol environment. The goal of this study was not only to predict energetically favorable binding sites of Mg^2+^ at the MOP receptor, but also to provide atomic details of the molecular mechanism regulating the positive allosteric modulation of the receptor induced by this divalent cation. This information is important considering that magnesium is known to enhance opioid-induced analgesia in different types of surgical procedures without increasing opioid side effects at therapeutic doses, thus suggesting that its use in combination with opioid analgesics may be useful in clinical practice (25).

## Methods

### MD simulations and system setup

Three different simulation setups were built for the ligand-free murine MOP receptor (residues 65-347), using the CHARMM-GUI web server (26). While one of them was based on the X-ray crystal structure of the inactive MOP receptor (PDB entry: 4DKL(27)), the other two system setups used the crystal structure of the active MOP receptor (PDB entry 5C1M (28)) and differed in the protonation state of the highly conserved D114^2.50^ residue. The missing loop between TM5 and TM6 in 4DKL was added using the Modeller software (29). Both ligand-free inactive and active MOP receptor models were embedded in a lipid bilayer with a POPC:cholesterol ≈ 9:1 ratio and an area of 7.5×7.5 nm^2^. The membrane and protein were then solvated with explicit TIP3P water molecules and 0.15 M concentrations of both Na^+^ and Mg^2+^ cations, and neutralized with a 0.3 M concentration of Cl^−^ ions. Each full simulation system contained approximately 70,000 atoms and had an initial volume of 7.5×7.5×11.9 nm^3^. The CHARMM36 force field (30) was used to model protein, lipids, and ions and all-atom MD simulations were carried out using the GROMACS software package version 2018.1 (31). Following a first energy minimization step, six short equilibrations runs with gradually decreasing harmonic constraints on lipid and protein heavy atoms were carried out, following the CHARMM-GUI membrane builder equilibration protocol. After these short equilibration runs, the three systems were further equilibrated for 100 ns in the NPT ensemble without constraints using the Nosé-Hoover thermostat (32) at 310 K (coupling constant *τ*_*t*_ = 1.0 ps) and the Berendsen barostat (33) at 1 atm (coupling constant *τ*_*p*_ = 5.0 ps). A final equilibration run of 20 ns was carried out under the production simulation conditions, in which the Berendsen barostat was replaced by the Parrinello-Rahman barostat (34) (coupling constant of *τ*_*p*_ = 5.0 ps at 1 atm). Different production runs ranging between ~0.7 μs and ~2.4 μs, for a total simulation time of over 20 μs, followed (see Table S1 for details). Long-range electrostatic interactions (for inter-particle distances beyond 1.2 nm) were taken into account using the Particle-Mesh Ewald (PME) algorithm (35). The van der Waals interactions were smoothly switched off gradually between 1.0 and 1.2 nm. Periodic boundary conditions were applied to the simulation boxes, and an integration time step of 2 fs was used for all simulations.

### Identification of cation binding sites

We used a graph theory-based approach to identify the preferred binding sites for each cation on each of the simulated MOP receptor systems. First, so-called cation binding *micro-sites* were detected by the presence of either Na^+^ or Mg^2+^ cations within the closest minimum distance (*r*_min_) from heavy atoms of at least two protein residues. The value of *r*_min_ for the two cations was determined by selecting the minima with the shortest distance in the radial distribution functions (RDFs) of the Na^+^-protein and Mg^2+^-protein distances, respectively, calculated from combined MD trajectories of all simulated active and inactive MOP receptor systems. Based on the RDF shown in Fig. S1, and in agreement with published work (18), the closest minimum distances were *r*_min_ = 2.9 Å and *r*_min_ = 4.8 Å for Na^+^ and Mg^2+^cations, respectively, suggesting that unlike Na^+^, Mg^2+^ binds to the protein through water molecules only. This is reasonable given that the Mg^2+^ hydration free-energy is known to be five times larger than the one of the Na^+^ cations (36). Given the large number of micro-sites calculated using the aforementioned criterion, for each simulated MOP receptor system, we aggregated micro-sites into the so-called *macro-sites* of cation binding using graph theory concepts. Specifically, for each simulated system, each micro-site represented a node in a graph with edges between nodes indicating common protein residues involved in cation coordination. A greedy inference algorithm for the stochastic block model (SBM) (37, 38) implemented in the Python package graph-tool (https://graph-tool.skewed.de/) was applied to identify the best connected sub-graphs within the larger graph, each of which corresponded to a macro-site. Fifty models with slightly different description length (DL) were generated and the model with the smallest DL was selected for the final partitioning of the micro-site graph into macro-sites.

### Calculation of cation binding affinity at macro-sites

The cation binding affinity 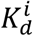 for the *i*-th macro-site was estimated based on its marginal occupancy probability *p*_*i*_, defined as the fraction of simulation frames in which the macro-site is occupied. Assuming that binding to the protein does not change the ion concentration, the binding affinity *K*_*d*_ can be estimated via the relation

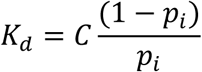

where *C* is the molar concentration of each cation species present in the simulation box. The lower and upper bound errors are estimated as 25^th^ and 75^th^ percentiles from the block average analysis of the combined trajectories for each system by dividing the trajectories into five equally sized blocks.

### Allosteric coupling between macro-sites of cation binding

We assessed the effect of each cation species on the binding of the other by calculating cooperativity coefficients between pairs of binding macro-sites. Specifically, we considered the joint probability distribution 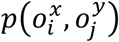 of the cation species *x* occupying the macro-site *i* and the cation species *y* occupying the micro-site *j*, marginalizing the occupancy of all the other macro-sites on the receptor. Indicating with 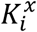 the affinity of the cation species *x* at macro-site *i* when macro-site *j* is not occupied, and correspondingly, with 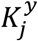 the affinity of the cation species *y* at macro-site *j* when macro-site *i* is not occupied, the presence of a cation at macro-site *j* changes the affinity of the cation species *x* at macro-site *i* to 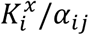, while the closure of the thermodynamic cycle ensures that when macro-site *i* is occupied, the cation binding affinity at macro-site *j* becomes 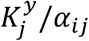. We report these cation binding affinity changes as log *α*_*ij*_, with negative values implying negative modulation, i.e. a reduction in the cation binding affinity at a given macro-site induced by the presence of another cation species. We estimate the values of *α*_*ij*_ from the ratios:

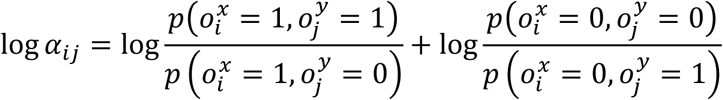

### Quantification of cation effect on extracellular loop gating

To elucidate the effect of Mg^2+^ binding on the conformational plasticity of the extracellular loop 2 and 3 (ECL2/3) region of the receptor, which has often been linked to the allosteric modulation of ligand binding, we outlined a simple activation model of the MOP receptor, and used our simulations to derive its thermodynamics properties. Specifically, our simple model only considers agonist binding and defines the active state of the receptor (with either neutral or charged D^2.50^) as the ligand-bound state, while the inactive receptor conformation is classified as a ligand-unbound state. We consider two different states of the ECL2/3 region of the MOP receptor, which we labeled open (O) or closed (C), depending on whether the minimum distance (*d*_Loop_) between the heavy atoms of residue pairs E310^ECL3^−R211^ECL2^ and Q212^ECL2^−D216^ECL2^ sampled during simulation was larger or smaller, than 10 Å, respectively. Indicating with *K*_*O*_ and *K*_*C*_ the ligand binding affinity for the ECL2/3 region open and closed states, respectively, and with *H*_*B*_ and *H*_*U*_ the equilibirium constants for ECL2/3 region opening in the bound and unbound receptors, we have that *K*_*C*_/*K*_*O*_ = *H*_*B*_/*H*_*U*_, and by applying the mass-action law, we can write the ligand bound fraction of the receptor at ligand concentration [*L*] as:

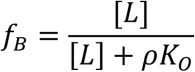

where

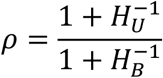

is a coefficient that measures the allosteric coupling between ligand binding and loop dynamics. If the dynamics of the ECL2/3 region does not depend on the presence of the ligand, then *H*_*U*_ = *H*_*B*_ and *ρ* = 1. In this case, the ligand binding thermodynamics is not affected by ECL2/3 region gating, and the ligand binding affinity is *L*_50_ = *K*_*O*_, i.e., the same as to the open receptor. If, however, the closed state of the ECL2/3 region is more stable in the ligand-bound complex, *H*_*U*_ ≪ *H*_*B*_ and *ρ* ≪ 1. In this case, *L*_50_ ≪ *K*_*O*_, i.e. the affinity of the ligand, and the stability of the active state, are increased by ECL2/3 region closure. Accordingly, this model can capture the basic mechanism by which ECL2/3 region closure increases the binding affinity of a ligand.

To quantify the effect of Na^+^ and Mg^2+^ cations on ECL2/3 region gating, we assessed the coupling between the ECL2/3 loop region conformational dynamics and cation binding. For a given Mg^2+^ concentration [*M*], we characterized the dynamics between the Mg^2+^-bound state of the open ECL2/3 region (*O*_*M*_) and the Mg^2+^-free state of the open ECL2/3 region (*O*) as *O*_*M*_(*i*) = *J*_*O*_(*i*)*O*(*i*)[*M*], where *J*_*O*_(*i*) is the equilibrium constant reflecting the affinity of Mg^2+^ for the open state, and *i* indicates the MOP receptor state (unbound/inactive, bound/active). Similarly, for the closed state of the ECL2/3 region, *C*_*M*_(*i*) = *J*_*C*_ (*i*)*C*(*i*)[*M*]. Indicating with 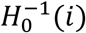 the equilibrium constant between the open and closed states of the ECL2/3 region in the absence of Mg^2+^, at magnesium concentration [*M*], we have that the equilibrium between the ECL2/3 region open and closed states is:

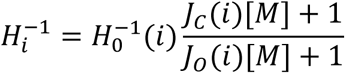

Using this expression for 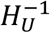 and 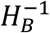 we obtain the dependence of the allosteric coefficient *ρ* on the Mg^2+^ concentration. Specifically, in the absence of magnesium, 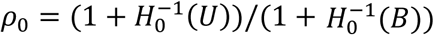, while for high magnesium concentrations 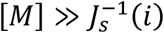, the allosteric coefficient is:

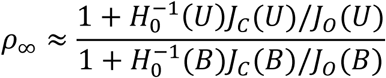

The values of *J*_*C*_ (*U*) and *J*_*O*_(*U*) (see Table S2) can be estimated from the occupancy of the extracellular binding sites in the closed and open states of the ECL2/3 region in the inactive MOP receptor trajectories, while the value of 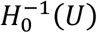 was obtained from the probability of the ECL2/3 region being in an open state in the same system. The values of *J*_*C*_ (*B*), *J*_*O*_(*B*), and 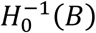 (see Table S2) were estimated from the corresponding fractions in the inactive and active MOP receptor trajectories. Values of *ρ*_0_ and *ρ*_∞_ are reported in Table S3.

### Calculation of co-information

To elucidate the molecular determinants of the positive allosteric modulation of the MOP receptor induced by the Mg^2+^ cation binding, we calculated the co-information between receptor residue pairs and Mg^2+^ occupancy of macro-sites within the ECL2/3 region of the MOP receptor. Specifically, residue dynamics was described using the Cartesian coordinates of the center of mass of the residue heavy atoms *R*_*i*_, while Mg^2+^ binding was described using a binary variable representing the occupancy state of any of the Mg^2+^ macro-sites in the ELC2/3 region, 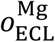. The mutual information *MI*(*R*_*i*_, *R*_*j*_) is a common quantity used to describe the long-range allosteric coupling between the dynamics of different regions of the receptor. In this work, we assessed the modulatory effect of Mg^2+^ binding by calculating the so-called co-information quantity:

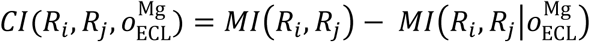

where *MI*(*R*_*i*_, *R*_*j*_) is the mutual information between the two residues, and 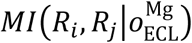 is the conditional mutual information between the residues, conditioned upon Mg^2+^ occupancy. This co-information is the difference between the information shared by *R*_*i*_ and *R*_*j*_ for a given value of 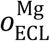 and the information shared by *R*_*i*_ and *R*_*j*_ irrespective of 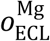, and thus measures the effect of 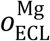 on the amount of information shared between *R*_*i*_ and *R*_*j*_. We note that negative values of the co-information indicate that 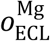 explains (some of) the observed correlation between *R*_*i*_ and *R*_*j*_.

Table S4 lists the normalized co-information values derived from:

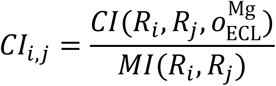

To reduce calculation time, trajectories were analyzed with a time step of 0.2 ns, and the mutual information and conditional mutual information values needed to calculate *CI*_*i*,*j*_ were estimated using the mdentropy python library (39) and the k-nearest neighbor algorithm with 5 bins per Cartesian dimension.

## Results and Discussion

To elucidate the atomic details of the mechanism by which Mg^2+^ binds and allosterically modulate the MOP receptor, we carried out MD simulations of experimentally-determined inactive and active structures of the receptor embedded in a lipid-water environment, and in the presence of both Na^+^ and Mg^2+^ cations. Unlike the inactive MOP receptor state, in which the highly conserved D^2.50^ residue is likely to be charged and bound to a Na^+^ cation, the D^2.50^ does not bind Na^+^ in the receptor’s active state and its protonation state depends on its proximity to cations (23, 40). Thus, we carried out MD simulations of one ligand-free inactive MOP receptor crystal structure corresponding to PDB code 4DKL (27), and two forms of the ligand-free active MOP receptor crystal structure corresponding to PDB code 5C1M (28), differing only in the protonation state of the D^2.50^ residue.

Simulations were run at equal concentrations (0.15 M) of Na^+^ and Mg^2+^. While the Na^+^ concentration is close to extracellular physiological conditions, the Mg^2+^ concentration is significantly higher than its normal value under the same conditions, but its use was necessary to observe binding events of this divalent cation. Notably, the positive allosteric modulation of ligand-free receptors by Mg^2+^ cations can only be observed experimentally at high Mg^2+^ concentrations (0.1 – 0.5 M) as recently demonstrated for the adenosine A_2A_ receptor (18).

### Mg^2+^ binds predominantly to the MOP receptor extracellular region and exhibits higher binding affinity for the active receptor conformation

In order to characterize likely binding sites of Mg^2+^ on the inactive or active MOP receptor (so-called macro-sites) at physiological concentrations of Na^+^, we used the graph theory-based approach described in Methods section. Briefly, so-called micro-sites of single or multiple pairs of residues that were found to be within a certain distance cutoff from either Na^+^ or Mg^2+^ cations were grouped into macro-sites of cation binding. Notably, this approach was able to successfully identify as a macro-site the set of residues that coordinate Na^+^ at the allosteric binding pocket revealed by crystal structures of inactive GPCRs, giving confidence in the analysis. The analysis of the occupancy probability of these macro-sites (see Methods) revealed a small number of binding regions with significant marginal occupancy probability.

Figures 1-3 report molecular and energetic details of all macro-sites identified with cation binding affinity less than 8 M in simulations of ligand-free inactive MOP receptor, ligand-free active MOP receptor with charged D^2.50^, and ligand-free active MOP receptor with neutral D^2.50^, respectively. Values of the Na^+^ and Mg^2+^ binding affinities for these macro-sites derived from occupancy probabilities (see Methods for details) are shown in these figures. As expected, in the ligand-free inactive MOP receptor system (Figure 1), the Na^+^ cation has the highest affinity for the allosteric binding site (in particular, residues D114^2.50^, N150^3.35^, S154^3.39^, N328^7.45^, and W293^6.48^) revealed by high-resolution crystal structures of various GPCRs. We calculate a *K*_*d*_ = 0.05 M for Na^+^ at this site (Figure 1e), which agrees with our previously published calculations from simulations of the MOP receptor in the presence of Na^+^ only, and their experimental validation (41). In addition to the crystallographic binding site, the Na^+^ cation was also found to bind at an extracellular site (specifically, residues N127^2.63^, D216^ECL2^, C217^ECL2^, T218^ECL2^; see Figure 1d) in the inactive MOP receptor system, albeit with the significantly lower affinity of 3.4 M. Unlike Na^+^, Mg^2+^ ions bound primarily at three slightly different sites on the extracellular region of the inactive MOP receptor system, which all had the D216^ECL2^ residue coordinating the cation (Figure 1a-c). The Mg^2+^ cation binding affinity for these sites was equally low, ranging from 3.6 M to 5.0 M. Not shown in this figure is a Mg^2+^ binding site observed in the vicinity of helix 8, which is likely an artifact of having run simulations with a zwitterionic C-terminus.

**Figure 1.**
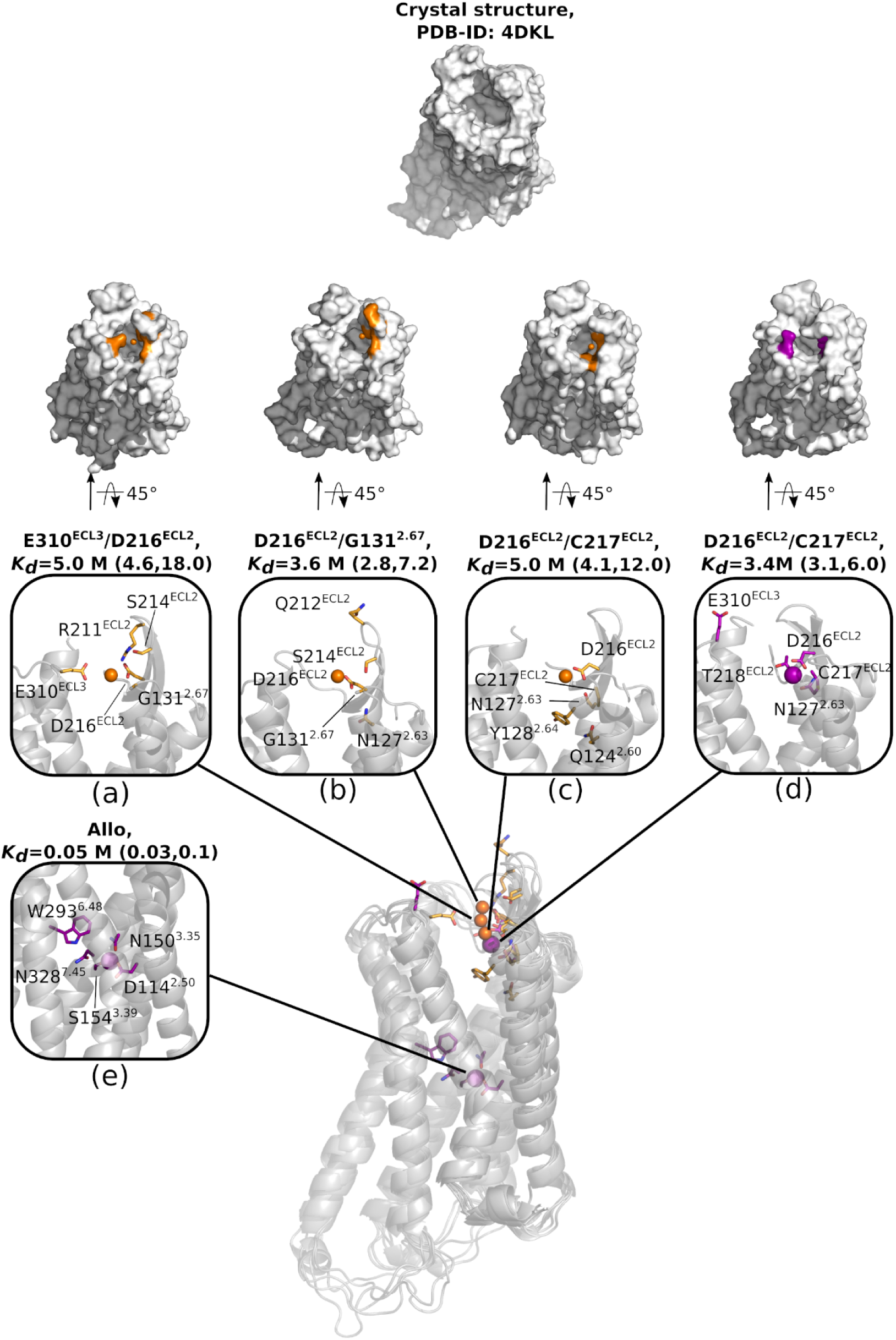
Molecular and energetic details of all macro-sites identified with cation binding affinity less than 8 M in simulations of ligand-free inactive MOP receptor. The top 5 representative residues of each cation binding macro-site (i.e., residues that are most frequently involved in Na^+^ and Mg^2+^ binding in micro-sites) are indicated in purple and orange colors, respectively, on the surface representations of the inactive MOP receptor. In the insets, the same residues are shown as sticks, while purple and orange spheres refer to Na^+^ and Mg^2+^ cations, respectively. Macro-sites in the extracellular region of the MOP receptor are labeled by the 2 residues that most frequently bind Na^+^ and Mg^2+^ cations in micro-sites.

While Na^+^ had a much higher affinity than Mg^2+^ for the inactive MOP receptor, this difference got reduced in the simulations of the active MOP receptor systems (see Figures 2 and 3). For instance, the affinity of Na^+^ for the crystallographic allosteric site decreased roughly 17-fold from about 0.05 M to 0.88 M in the ligand-free active MOP receptor system with a charged D^2.50^ (Figure 2d). This is consistent with the knowledge that this site is partially collapsed in active GPCR structures and not suitable to the same type of cation binding (19). The Na^+^ affinity for the extracellular region also decreased, albeit less dramatically, from 3.4 M to 4.9 M (Figure 2b). On the other hand, not only did the Mg^2+^ binding affinity for the extracellular loop regions slightly increase in the ligand-free active MOP receptor system with a charged D^2.50^ compared to the simulated inactive receptor system (from 3.6 – 5.0 M to 2.2 – 2.6 M; compare Figures 1a-c with Figures 2a and 2c), but the Mg^2+^ cation was also found to bind at the orthosteric ligand binding site defined by residue D147^3.32^ with a *K*_*d*_ of 0.7 M, the highest affinity value for a macro-site for any ion in this system (see Figure 2e). Notably, although the orthosteric ligand binding site was equally accessible in the ligand-free inactive MOP receptor system, the Mg^2+^ cation did not bind at this site during simulation of the ligand-free inactive MOP receptor system.

**Figure 2.**
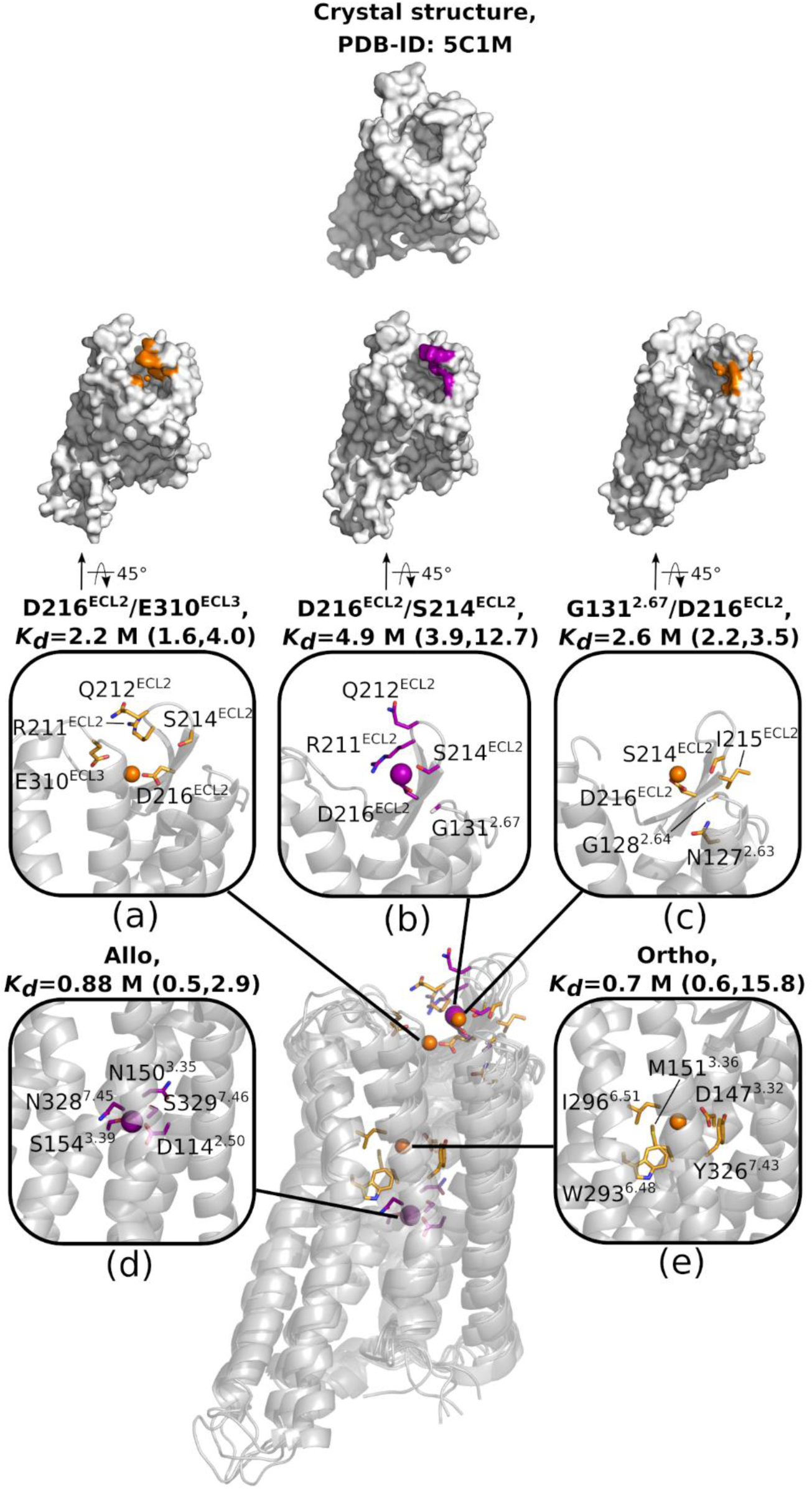
Molecular and energetic details of all macro-sites identified with cation binding affinity less than 8 M in simulations of the ligand-free active MOP receptor with a charged D^2.50^ residue. See caption of Figure 1 for details.

**Figure 3.**
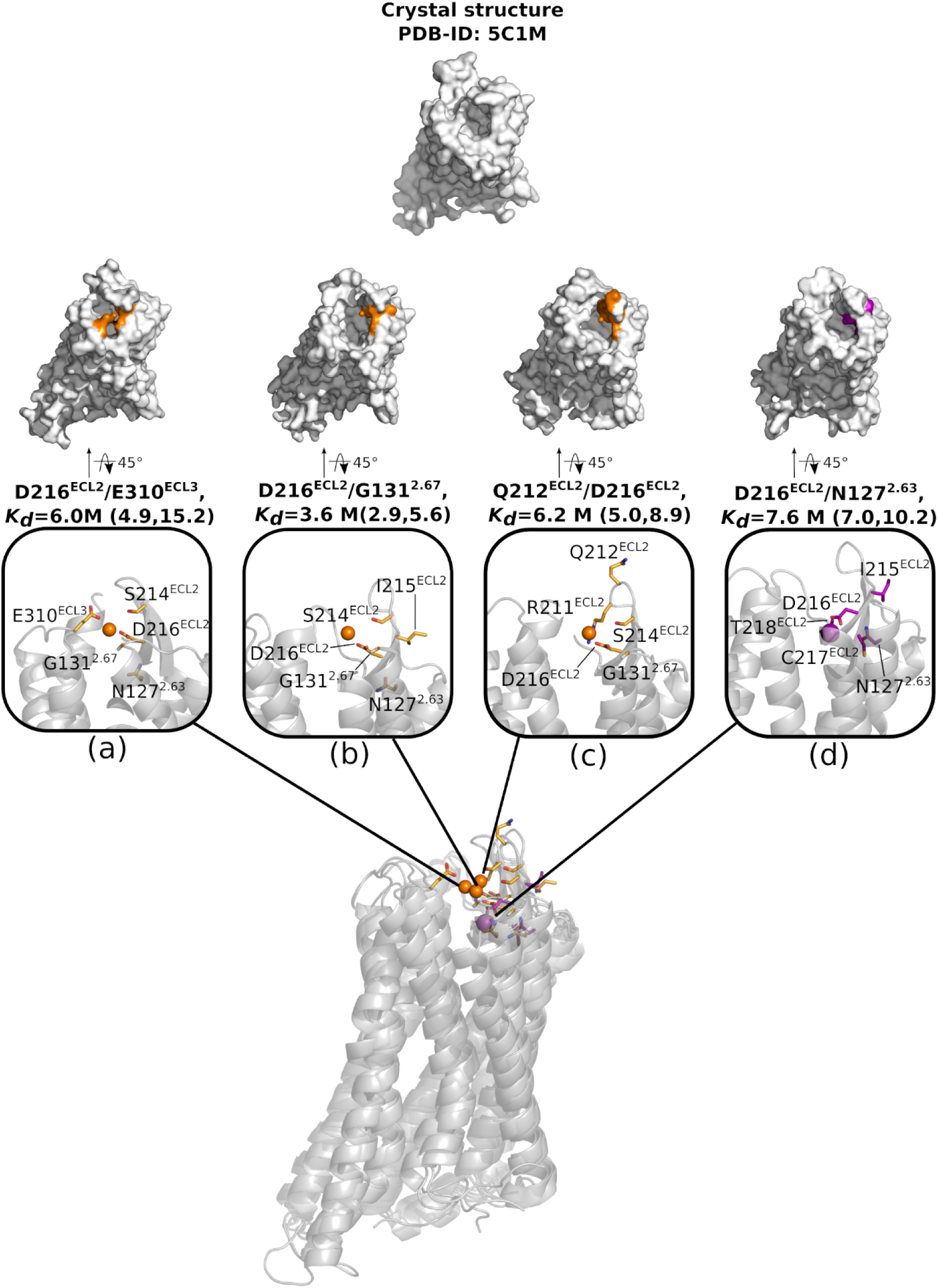
Molecular and energetic details of all macro-sites identified with cation binding affinity less than 8 M in simulations of the ligand-free active MOP receptor with a neutral D^2.50^ residue. See caption of Figure 1 for details.

Intriguingly, the change of the D^2.50^ protonation state to neutral in the simulated active MOP receptor system had a significant impact on the binding affinity of both Na^+^ and Mg^2+^ cations for the ligand-free MOP receptor (Figure 3). In this system, both the Na^+^ affinity for the allosteric binding site defined by D^2.50^ and the Mg^2+^ affinity for the orthosteric ligand binding site defined by D147^3.32^ fell below the cutoff value of 8 M chosen for significance. Notably, the affinity values of Mg^2+^ for the extracellular loop region (ranging from 3.6 to 6.2 M; see Figures 3a-c) were closer to those recorded for the ligand-free inactive MOP receptor system than for the active receptor system with a charged D^2.50^, whereas Na^+^ binding affinity for the extracellular region dropped to 7.6 M (Figure 3d).

### Mg^2+^ binding to the extracellular region of the MOP receptor promotes occlusion of the ligand binding pocket

A unique feature of Mg^2+^ binding to the MOP receptor extracellular region, particularly negatively charged residues D216^ECL2^ and E310^ECL3^, is that it can produce a transient closure of the orthosteric ligand binding pocket by bringing ECL2 and ECL3 closer together. These two extracellular loops effectively act as a “gate”, and when closer together, they can hinder access of the ligand to the pocket, as well as its departure from it. This phenomenon appears to be a unique feature of the divalent Mg^2+^ cation since it was not observed when the monovalent Na^+^ cation was bound at this site. The surface representations of the MOP receptor systems in Figures 1-3 clearly show the different degree of accessibility of the MOP receptor orthosteric ligand binding pocket in the presence of bound Na^+^ or Mg^2+^ in the extracellular region. Notably, visual inspection of the trajectories, as well as smaller values of the equilibrium constant *H*_*M*_ (see Table S2), reveal that the Mg^2+^-induced ECL2/3 loop closure occurs with higher frequency in the simulated active MOP receptor conformations than in the inactive receptor system, suggesting a coupling between activation and ECL2/3 closure.

To quantify the effect of Mg^2+^-induced ECL2/3 closure on ligand binding and receptor activation, we calculated a coupling coefficient, as normally done to measure the allosteric modulation between different receptor degrees of freedom. Specifically, we considered a first approximation in which the agonist binding affinity is expressed as *L*_50_ = *ρK*_*O*_, where *K*_*O*_is the agonist binding affinity to the receptor when the ECL2/3 gate is “open”, and *ρ* is an “allosteric coefficient” that takes into account the Mg^2+^occupancy of the extracellular binding sites in the closed and open states of the ECL2/3 region, thus capturing the effect of cation binding on ECL2/3 closure. The same coefficient *ρ* was used to express the constitutive activity ratio between active and inactive MOP receptors as *R*/*ρ*, where *R* is the active/inactive receptor ratio in the absence of ECL2/3 gating. Values of the *ρ* coefficient calculated for different Mg^2+^ concentrations are shown in Table S3 (see Table S2 for values of its components and Methods for equations). In the absence of allosteric modulation by the cation, *ρ* is expected to be equal to 1. On the other hand, if *ρ* < 1, the agonist binding affinity is predicted to increase as is the active fraction of the receptor. Comparison between the inactive MOP receptor system and the active MOP receptor with neutral D^2.50^ yields *ρ*(high Mg^2+)^/*ρ*(no Mg^2+^) ≈ 0.25, which suggests a 4-fold increase in agonist binding affinity in the presence of magnesium, as well as a 4-fold increase of the constitutively active fraction of the receptor (see Table S3). Similarly, comparing the inactive MOP receptor and the active MOP receptor with charged D^2.50^ yields a value of *ρ*(high Mg^2+)^/*ρ*(no Mg^2+^) ≈ 0.20, corresponding to a 5-fold change (see Table S3).

### Na^+^ and Mg^2+^ compete for binding at ECL2 sites

To address possible binding competition between Na^+^ and Mg^2+^ cations for specific sites on the MOP receptor, we calculated the allosteric modulation coefficient log *α*(*i*_*Na*_, *j*_*Mg*_) for the cation binding affinity for each of the three simulated MOP receptor systems. Negative values of this coefficient in the plots of Figure 4 (red color) indicate a negative cooperativity, i.e. that the binding of Na^+^ at site *i*_*Na*_ decreases the affinity of Mg^2+^ at site *j*_*Mg*_ (and vice versa), while positive values (in blue in Figure 4) would suggest that binding of one ion increases the affinity of the other. Notably, most of the coupling coefficients in the plots of Figure 4 are negative, suggesting that there is active competition for the available binding sites in all simulated MOP receptor systems. Specifically, in the inactive MOP receptor system (Figure 4a), Mg^2+^ strongly competes (large negative values of log *α*) with Na^+^ for any of the extracellular binding sites. Notably, the coupling between the Na^+^ allosteric binding site and the Mg^2+^ E310^ELC3^/D216^ECL2^ site is also negative, albeit not very pronounced (log *α* ≈ −1.0), suggesting a 3-fold reduction of the Na^+^ affinity for its allosteric binding site in the presence of Mg^2+^ at the E310^ELC3^/D216^ECL2^ site in the inactive MOP receptor.

**Figure 4:**
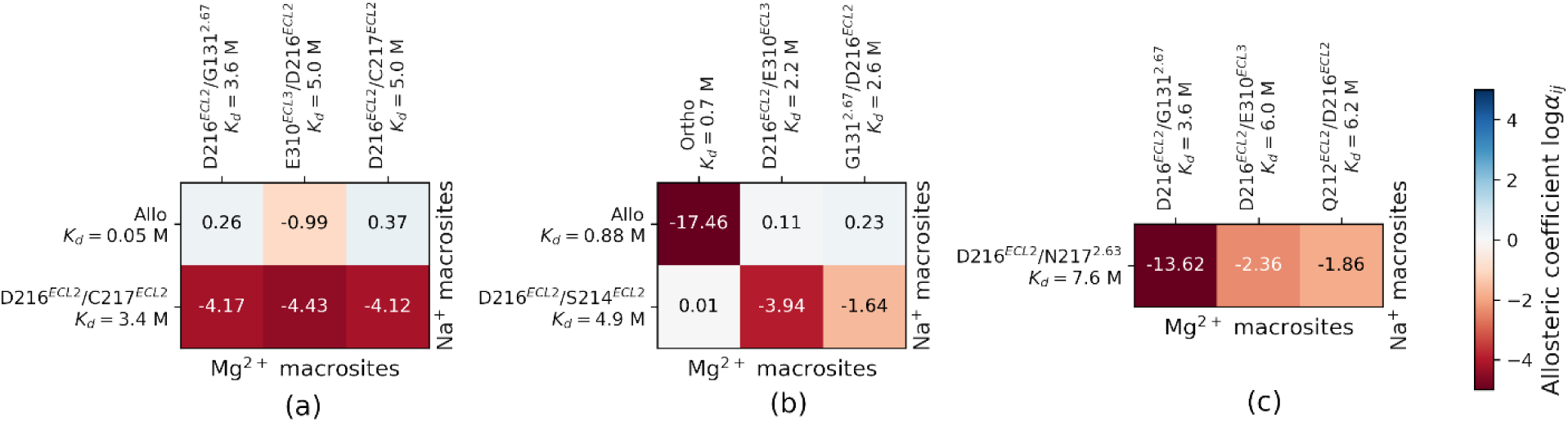
Allosteric coefficients for cooperative binding at Na^+^ and Mg^2+^ binding sites in the simulated MOP receptor systems. Panels a, b, and c, refer to inactive, active with charged D^2.50^, and active with neutral D^2.50^ MOP receptor, respectively. The coefficients are reported as log *α*, with negative values (in red) corresponding to negative cooperativity, and positive values (blue) corresponding to positive cooperativity of the cation binding affinity.

In the active MOP receptor (Figures 4b and 4c), a negative coupling is also observed between all pairs of binding sites in the extracellular region of the receptor. Furthermore, in the active receptor with a charged D^2.50^ (Figure 4b), we observe a remarkable negative modulation (log *α* ≈ −17.0) of the binding of Mg^2+^ to the orthosteric ligand binding pocket induced by Na^+^ binding at the D^2.50^ allosteric binding site. This high negative cooperativity appears to be the reason for the lack of Mg^2+^ binding at the orthosteric ligand binding site when a Na^+^ cation is present at its allosteric binding site, and vice versa.

### Cation binding modulates the information flow across the receptor

In order to further elucidate the molecular mechanism underlying Mg^2+^ positive allosteric modulation of the MOP receptor, we analyzed the impact of Mg^2+^ binding on the information flow across the receptor. Reasoning that receptor allosteric modulation can be captured by the mutual information between residues, we characterized the residues mediating the modulatory effect of the cation by calculating the normalized co-information between pairs of receptor residues (*i,j*) and Mg^2+^ occupancy of macro-sites within the ECL2/3 region of the MOP receptor (see Methods section).

The receptor residue pairs whose correlation is maximally affected by Mg^2+^ binding (i.e., the top 20 pairs with the highest co-information values) are listed in Table S4 and their location shown in Figure 5 for the three simulated MOP receptor systems. Accordingly, in the inactive MOP receptor system (Figure 5a), Mg^2+^ binding to the ECL2/3 region most dramatically affects the mutual information between specific residues in ECL2 (i.e., Y210, R211, I215, and S222), ECL3 (i.e., P309), TM2 (i.e., A102^2.38^, N109^2.45^, A113^2.49^), TM3 (D147^3.32^, Y149^3.34^, N150^3.35^, M151^3.36^, and T153^3.38^), ICL2 (i.e., L176), TM5 (i.e., C235^5.41^), TM6 (i.e., V288^6.43^ and C292^6.47^), the intracellular end of TM7, flanking the NPxxY motif (L324^7.41^, C330^7.47^, L331^7.48^, L339^7.56^), ICL4 (i.e., E341), and H8 (i.e., N342 and K344).

**Figure 5:**
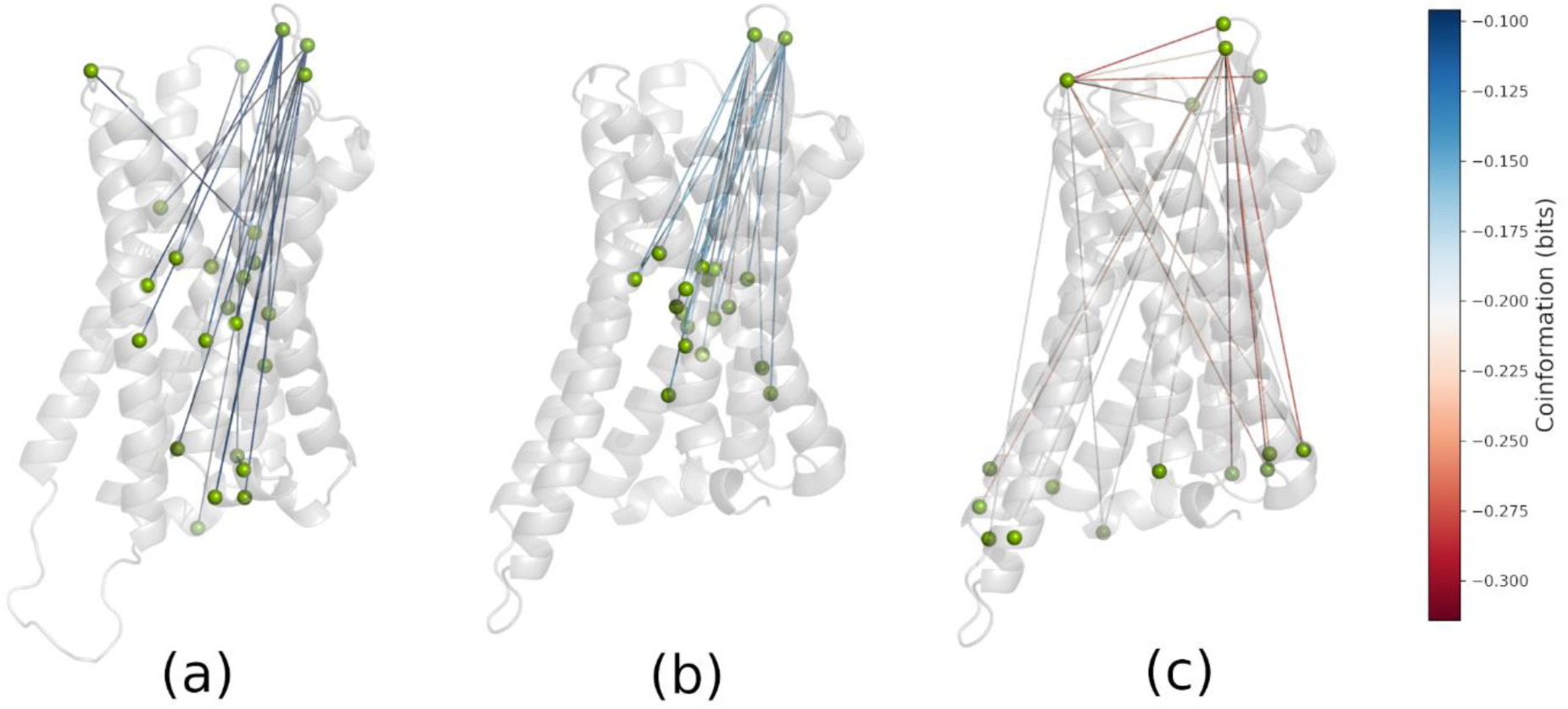
Location of the MOP receptor residues pairs whose correlation is maximally affected by Mg^2+^ binding in the three simulated receptor systems. Panels a, b, and c, refer to inactive, active with charged D^2.50^, and active with neutral D^2.50^ MOP receptor, respectively. Lines between residues are colored according to their co-information value, with the most negative values shown in red.

Notably, co-information values are much more negative in the simulated active MOP receptor systems, and especially in the system with a neutral D^2.50^ (see Table S4 and Figure 5c), reflecting a tighter coupling between magnesium occupancy of macro-sites within the ECL2/3 loop region and the receptor dynamics in the active conformational state. In the active MOP receptor with a charged D^2.50^ residue, the most affected regions of the receptor by Mg^2+^ binding share many similarities with those of the inactive MOP receptor (compare Figures 5a and 5b), involving specific residues within ECL2 (Y210^ECL2^ and R211^ECL2^), TM2 (N109^2.45^), TM3 (N150^3.35^, M151^3.36^, and T153^3.38^), TM6 (C292^6.47^), and TM7 (L324^7.41^ and L331^7.48^). On the other hand, the co-information observed in the active receptor with neutral D^2.50^ is strikingly different from that of the MOP receptor with charged D^2.50^ (compare Figures 5b and 5c). Notably, among the most affected receptor regions by cation binding in this system are ECL2/3 residue pairs, specifically: P309^ECL3^ with K209^ECL2^, R211^ECL2^, Q212^ECL2^, or S222^ECL2^, as well as the intra-ELC2 pair R211^ECL2^-S222^ECL2^. As expected by a positive allosteric modulator, magnesium binding also modulates the information transfer from ECL2/3 residues (i.e., R211^ECL2^, S222^ECL2^, and P309^ECL3^) to the intracellular G protein-binding region of the MOP receptor, including the intracellular ends of TM1 (I93^1.57^ and V94^1.58^), TM3 (C170^3.55^), TM4 (P181^4.39^), TM5 (L257^5.63^ and K260^5.66^), and TM6 (D272^6.27^ and R273^6.28^), as well as ICL1 (M99), ICL2 (V173), and ICL4 (D340).

Taken together, these results support a mechanism by which Mg^2+^ binding to the MOP receptor active state with neutral D^2.50^ (i.e., the expected most probable protonation state for an active receptor) has a direct effect on the dynamics of the ECL2/3 region, which in turn is strongly coupled with the dynamics of the intracellular region of the receptor, whereas this coupling is less effective in the inactive receptor. Notably, a similar mechanism was recently proposed for a different GPCR subtype (18), suggesting a possible molecular paradigm of GPCR allosteric modulation by cations.

## CONCLUSIONS

Our simulations provide atomic details of the molecular mechanism by which magnesium cations bind to the extracellular region of the MOP receptor and allosterically modulate the G-protein-binding region of the receptor. Specifically, they support a mechanism by which Mg^2+^ cations, unlike Na^+^ cations, promote states of the active MOP receptor with ECL2 and ECL3 folding back over the extracellular opening of the receptor TM bundle, thus impairing agonist unbinding, and possibly enhancing agonist binding affinity. Mutations of residues involved in Mg^2+^ binding to the ECL2/3 region (e.g., the acidic residues E310^ELC3^ and D216^ECL2^), as well as mutations of residues identified as mediators of the cation allosteric effect (e.g., R211^ECL2^, Q212^ECL2^, and P309^ECL3^) are predicted to affect its positive allosteric modulation of the MOP receptor.

While Na^+^ and Mg^2+^ cations actively compete for binding to extracellular sites of the MOP receptor, a larger number of binding sites is available to Mg^2+^ in this region. Thus, despite the low (molar) range of affinities for these sites, we would predict that Mg^2+^ cations bind to the extracellular region of the MOP receptor even at lower concentrations. Notably, Mg^2+^ was not observed to bind to the Na^+^ allosteric site, most likely because of its double positive charge and because of an unfavorable balance between desolvation and binding. Mg^2+^ could, however, bind at the orthosteric ligand binding site with a significantly higher affinity than sodium. Furthermore, we observed a strong negative cooperativity between Mg^2+^ binding at the orthosteric site and Na^+^ binding at the allosteric site, suggesting that Mg^2+^ binding could reduce the fraction of bound Na^+^, thus reducing its inhibitory effect.

## Author Contributions

X.H. carried out the simulations. All authors analyzed the results and wrote the manuscript.

## Acknowledgments

This work was supported by National Institutes of Health grant DA045473. Computations were run on resources available through the Scientific Computing Facility at the Icahn School of Medicine at Mount Sinai and the Extreme Science and Engineering Discovery Environment under MCB080077, which is supported by National Science Foundation grant number ACI-1053575.

